## Supplementary material for "ImmuneBuilder: Deep-Learning models for predicting the structures of immune proteins": Supplemetary information

### Appendix A NanoBodyBuilder2

Nanobodies are single domain antibodies found in organisms such as camelids and sharks [1]. Their structure is similar to that of an antibody heavy chain variable region. However, the CDR-H3 loop in antibodies and nanobodies often take on very different conformations. While in antibodies the number of possible CDR-H3 conformations is limited by the light chain [2], the lack of light chain in nanobodies allows them to adopt a wider range of conformations [3]. To not bias the model towards light chain constrained CDR loop conformations, we trained separate models for antibody and nanobody structure prediction.

#### A.1 Methods

The model architecture, training procedure, model selection and structural refinement for NanoBodyBuilder2 is the same as for ABodyBuilder2 and is described in the main text. To train NanoBodyBuilder, 1935 nanobody structures were extracted from SAbDab [4] on the 10th of July 2022, fifty of which were randomly selected as a validation set. For the benchmark data, forty structures released between January and July 2022, with a resolution better than 2.3Å and resolved via X-ray diffraction were selected. It was ensured that there were no structures with the same sequence in the train, tests and validation sets. A full list of these structures is given at https://github.com/oxpig/ImmuneBuilder.

#### A.2 Results

In this section we compare NanoBodyBuilder2, our nanobody-specific method, to three other tools used for nanobody structure prediction. We compared against two homology modelling methods (the original version of ABodyBuilder [5] and MOE [6]), and one general protein structure prediction method (AlphaFold2 [7]). As a benchmark, we selected a non-redundant set of forty nanobody structures recently added to SAbDAb [4]. None of the benchmark structures are in the training or validation set for either of the deep learning methods used.

In Table A1, we report results on the same metrics as were used for antibodies in the main text. The exceptions to this are metrics involving the light chain, such as heavy and light chain packing angles or light chain regional RMSDs.

| Method | CDR1 | CDR2 | CDR3 | Fw |
|---|---|---|---|---|
| ABodyBuilder | 3.12 | 2.19 | 5.29 | 1.13 |
| MOE | 2.67 | 1.99 | 4.90 | 1.19 |
| AlphaFold2 | 2.08 | **1.35** | 3.44 | 0.82 |
| NanoBodyBuilder2 | **1.98** | 1.37 | **2.89** | **0.79** |

| Method | $\chi 1$ | $\chi 2$ | $\chi 3$ | $\chi 4$ | E/B |
|---|---|---|---|---|---|
| ABodyBuilder | 0.72 | **0.74** | **0.54** | **0.63** | 0.92 |
| MOE | 0.68 | 0.63 | 0.43 | 0.49 | 0.92 |
| AlphaFold2 | **0.78** | **0.74** | 0.51 | **0.63** | **0.94** |
| NanoBodyBuilder2 | 0.77 | 0.70 | 0.53 | 0.58 | 0.93 |

| Method | Peptide bond | Clash | D-amino acid | Cis-bond |
|---|---|---|---|---|
| ABodyBuilder | 49 | 0 | 8 | 21 |
| MOE | 0 | 7 | 6 | 0 |
| AlphaFold2 | 0 | 0 | 0 | 0 |
| NanoBodyBuilder2 | 0 | 0 | 0 | 0 |

**Table A1** Comprehensive benchmark between ABodyBuilder, MOE, AlphaFold2 and NanoBodyBuilder2 for predicting nanobody structures. In the first table, the mean RMSD to the crystal structure across the test set for each of the three CDRs and the framework is shown. In the second, we show the accuracy at modelling each of the first four torsion angles of the side chain ($\chi$) and the accuracy at predicting whether a residue is exposed or buried (E/B). The third table shows the total number of the stereochemical errors found in the predicted structures. For a more a more in depth description of each metric, see the results section of the main text.

NanoBodyBuilder2 predicts the backbone structure of nanobodies with higher accuracy than all other benchmarked methods. The biggest improvement is in the CDR3 loop, which all methods model least accurately. NanoBodyBuilder2 also predicts the chi ($\chi$) side chain angles with an accuracy comparable to AlphaFold2 and the original version of ABodyBuilder. In terms of speed, AlphaFold2 takes around one hundred times longer to generate a nanobody structure than any of the other methods.

In Figure A1, the structure of an antibody heavy chain as predicted by ABodyBuilder2 is compared to the same chain as predicted by NanoBody-Builder2. Although both models predict a similar structure for the framework, the predicted CDR-H3 conformation is very different.

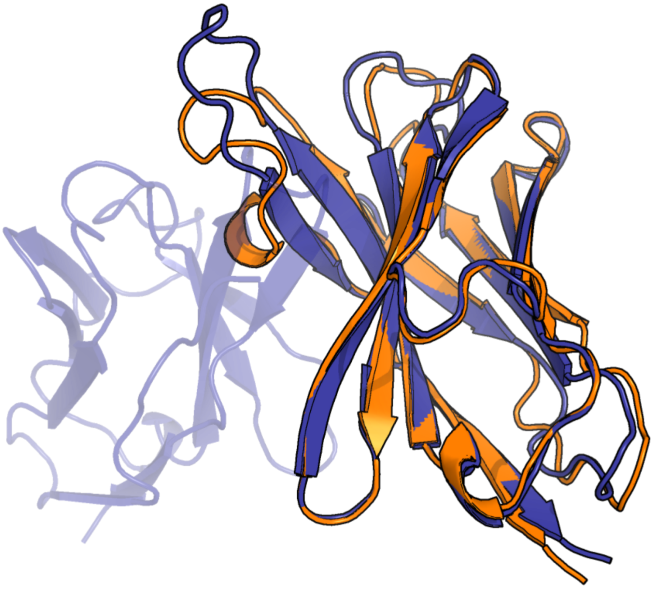

**Fig. A1** Example of an antibody heavy chain as predicted by ABodyBuilder2 and NanoBodyBuilder2. The NanoBodyBuilder2 prediction is shown in orange and the ABody-Builder2 prediction is shown in blue. While ABodyBuilder2 predicts the CDR-H3 to be in an erect conformation to avoid collisions with the light chain, NanoBodyBuilder predicts the loop to take a less compact form.

### Appendix B TCRBuilder2

The global structure of the variable domain in TCRs is similar to the antibody variable domain. However, antibody and TCR CDR loops tend to be of different lengths and mostly occupy distinct structural spaces [8]. As with nanobodies, we built a model for TCR structure prediction trained only on TCRs (TCRBuilder2).

#### B.1 Methods

The model architecture, training procedure, model selection and structural refinement for TCRBuilder2 is the same as for ABodyBuilder2 which is described in the main text. To train TCRBuilder2, the structures of 704 TCR variable domains were extracted from STCRDab [9] on 20th of May 2022, fifty of which were randomly selected as a validation set. For the benchmark data, the structures of 21 alpha-beta TCRs with a resolution better than 3.5Å and released between January and August 2022 were selected. It was ensured that there were no structures with the same sequence in the train, test and validation sets.

#### B.2 Results

In this section we compare TCRBuilder2 against two other methods for TCR structure prediction. We benchmarked it against a homology modelling method (the original version of TCRBuilder [10]) and a general protein structure prediction method (AlphaFold-Multimer [11]). Additionally, we compared it to our antibody-specific method (ABodyBuilder2) to showcase the benefits of training only on TCRs.

| Method | CDR-A1 | CDR-A2 | CDR-A3 | Fw-A | CDR-B1 | CDR-B2 | CDR-B3 | Fw-B |
|---|---|---|---|---|---|---|---|---|
| TCRBuilder | 1.60 | 1.31 | 2.89 | 0.87 | 0.99 | 0.90 | 3.12 | 0.81 |
| AlphaFold-M | **1.25** | 0.96 | **1.84** | **0.69** | 0.75 | 0.65 | 1.94 | 0.82 |
| ABodyBuilder2 | 3.49 | 6.57 | 3.14 | 2.89 | 3.27 | 3.77 | 3.48 | 3.65 |
| TCRBuilder2 | 1.34 | **0.93** | 1.85 | 0.90 | **0.74** | **0.63** | **1.93** | **0.67** |

| Method | HL | HC1 | LC1 | HC2 | LC2 | dc |
|---|---|---|---|---|---|---|
| TCRBuilder | 4.80 | 3.07 | 2.17 | 4.03 | 1.77 | 0.32 |
| AlphaFold-M | **2.95** | **1.45** | **1.97** | 2.83 | **1.51** | 0.36 |
| ABodyBuilder2 | 74.26 | 25.65 | 20.71 | 63.19 | 27.42 | 1.91 |
| TCRBuilder2 | 3.32 | 1.88 | 2.80 | **2.44** | 1.94 | **0.28** |

| Method | $\chi 1$ | $\chi 2$ | $\chi 3$ | $\chi 4$ | E/B |
|---|---|---|---|---|---|
| TCRBuilder | 0.70 | **0.69** | **0.53** | **0.53** | 0.90 |
| AlphaFold-M | **0.77** | **0.69** | 0.50 | 0.52 | **0.92** |
| ABodyBuilder2 | 0.58 | 0.52 | 0.36 | 0.39 | 0.83 |
| TCRBuilder2 | 0.76 | 0.67 | 0.50 | 0.47 | 0.91 |

| Method | Peptide bond | Clash | D-amino acid | Cis-bond |
|---|---|---|---|---|
| TCRBuilder | 74 | 29 | 0 | 6 |
| AlphaFold-M | 0 | 0 | 0 | 0 |
| ABodyBuilder2 | 0 | 0 | 0 | 0 |
| TCRBuilder2 | 0 | 0 | 0 | 0 |

**Table B2** Comprehensive benchmark between ABodyBuilder2, TCRBuilder, AlphaFold-Multimer and TCRBuilder2 for predicting TCR structures. The top table shows the mean RMSD to the crystal structure across the test set for each of the six CDRs and frameworks. The CDRs and framework (Fw) regions are labelled A for the alpha chain and B for the beta chain. The second table shows the mean absolute error in each of the six ABangles [12]. In the third table, the accuracy at modelling each of the first four torsion angles of the side chain ($\chi$) and the accuracy at predicting whether a residue is exposed or buried (E/B) is shown. The fourth table shows the total number of the stereochemical errors found in the predicted structures. For a more a more in depth description of each metric, see the results section of the main text.

ABodyBuilder2 fails to accurately model TCR structures, with the biggest errors being the modelling of the relative orientation between the alpha and beta chains. Both TCRBuilder2 and AlphaFold-Multimer predict structures with a comparable accuracy, outperforming the original version of

TCRBuilder. In terms of run time, TCRBuilder2 takes about as long as ABodyBuilder2 and TCRBuilder, which are all over a hundred times faster than AlphaFold-Multimer.

### Appendix C Heavy and light chain packing characterisation

To benchmark the accuracy of different methods at predicting the relative orientation between the VH and VL chains we use the absolute error in six values, taken from ABangle [12], that have been shown to fully characterise it. In Figure C2, we provide a brief description on how each ABangle is defined. For a more in depth description, please see the original paper [12].

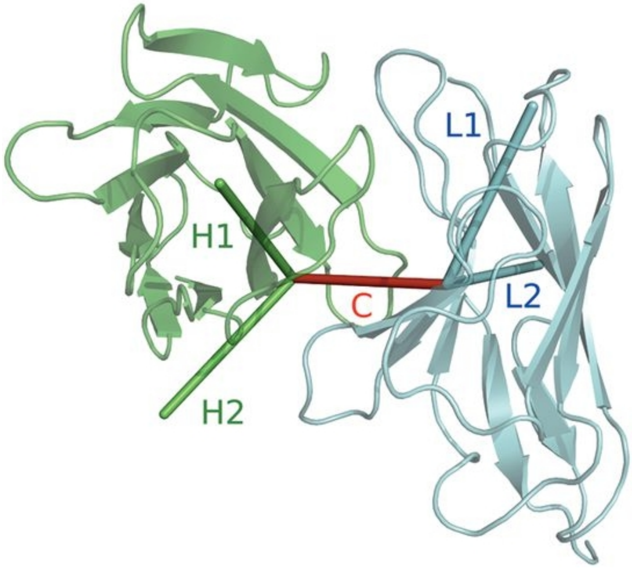


**Fig. C2** Antibody structure with the five vectors used to calculate the 6 values used to define heavy and light chain packing. The distance dc is defined as the length of the vector C. The HL angle is defined as the torsion angle between the H1 and L1 vectors measured about C. The HC1, HC2, LC1 and LC2 angles are the angles between the H1, H2, L1 and L2 vectors and the vector C. Figure taken from [12].
